# Association of miR-181a-5p with *Lantana camara* leaf extract-mediated inhibition of proliferation, survival, and migration in luminal A-type MCF-7 cells and triple-negative type MDA-MB-231 cells

**DOI:** 10.1101/2025.10.14.682348

**Authors:** Arundhaty Pal, Sourav Sanyal, Tapas Kumar Sengupta

## Abstract

**Ethnopharmacological relevance:** Breast cancer has a high mortality with increasing chemoresistance and recurrence. *Lantana camara* has diverse medicinal properties, including anti-cancer potential; however, its role in breast cancer is not well-explored.

**Aim of the study:** The study aims to elucidate the role of *Lantana camara* leaf extract in miR-181a-5p-mediated control of proliferation, survival, and migration of breast cancer cells.

**Material and methods:** Real-time PCR was employed for RNA expression studies. Bioinformatic tools were used for identifying miR-181a-5p target mRNAs and seed regions in the 3 prime untranslated region of target mRNAs. miR-181a-5p was overexpressed in cells using transfection method. Flow cytometry was employed for cell cycle and apoptosis assays. Scratch assay was performed to study cell migration. The 3 prime untranslated region of *CDKN3* was cloned, and site-directed mutagenesis was employed to mutate miR-181a-5p seed region. Western blot analysis was employed for a reporter assay.

**Results:** *L. camara* leaf extract upregulated the expression of miR-181a-5p with concomitant decrease in mRNAs of *BCL-2*, *MCL-1*, and *CDKN3* in breast cancer cells. Overexpression of miR-181a-5p induced cell death in MCF-7 and MDA-MB-231 cells, and cell death increased in the presence of leaf extract. miR-181a impeded migration in both cell lines in the presence of the extract. Reporter assay confirmed the interaction of miR-181a-5p with 3 prime untranslated region of the *CDKN3* mRNA.

**Conclusion:** Our study suggests that the extract may play an important role in regulating miR-181a-5p in MCF-7 and MDA-MB-231 cells, and may have important therapeutic potential for breast cancer therapy.

## 1. Introduction

Breast cancer is one of the most commonly reported cancers in females worldwide, and it also accounts for the majority of cancer-related deaths in females. Several factors like genetics, physiological condition, lifestyle habits, Smoking, and alcohol consumption may increase the risk of breast cancer. Women in their 50s are more prone to developing breast cancer; however, it can occur at any age. Depending on the severity of the breast cancer, it is subdivided into different subtypes-luminal A, luminal B, HER2–enriched, and triple-negative breast cancer (TNBC)(Al-thoubaity, 2020). Among different types of breast cancer, luminal A type contributes to almost 58.5% of the breast cancer burden, whereas TNBC accounts for 16% of all cases(Al-thoubaity, 2020). Luminal A-type breast cancer is slow growing, has a low grade, and has the best prognosis. In contrast, TNBC is the most aggressive, highly proliferative, and has high mutation rates in DNA repair genes, resulting in high genomic instability. TNBC has a very poor prognosis and a high relapse rate(Orrantia-Borunda et al., 2022). There are different treatment strategies for breast cancer depending on molecular subtypes, e.g., surgery, chemotherapy, radiation therapy, hormonal therapy, and targeted therapy. For treating luminal A-type breast cancer, drugs like tamoxifen and fulvestrant are used (Xiong et al., 2025). For TNBC, chemotherapeutic drugs like doxorubicin and taxanes are being used. Despite the success, chemotherapeutic agents have several limitations, like toxicity, drug resistance, varying patient outcomes, etc. Moreover, TNBC has higher tumor heterogeneity, a high relapse rate. Targeted therapies are also limited for TNBC (Obidiro et al., 2023). To overcome the limitations of conventional therapies, the discovery of more effective and safe treatment options is the utmost necessity. The plant world is a huge source of bioactive compounds, and only 10% of it is known till now (Iqbal et al., 2017). Different plant parts, such as stems, bark, leaves, flowers, and roots, are sources of metabolites like terpenoids, flavonoids, glycosides, saponins, and lignans. These compounds have the potential to act like anti-tumor agents. Some examples include: taxols, genistein, berberine, curcumin, etc (Iqbal et al., 2017).

*Lantana camara* is a flowering ornamental plant that belongs to the Verbenaceae family, also known as red or wild sage. Its nomenclature is verified via the World Flora Online (WFO) Plant List (https://wfoplantlist.org/). This plant was first introduced in the 19^th^ century in India (Wagh et al., 2023). There are 650 different types of plants found in over 60 other countries. Different parts of the plants, including flowers, leaves, and roots, have been used in traditional medicine to treat various ailments like fever, pox, flu, stomach ache, wounds, etc. Lantadene – A, B, C, and oleanolic acid isolated from the plant have attracted attention in drug research for their anti-cancer, antibacterial, AIDS, and anti-inflammatory activities (major) (Wagh et al., 2023).In various studies, including our own, it has been reported that extracts from the *Lantana* plant exhibit anti-tumor effects on different cancer cell lines, including those of breast cancer. Studies by Han et al. demonstrated that the ethanolic extract of *L. camara* could induce apoptosis in MCF-7 cells (Han et al., 2015). Another report by Shamsee et al. showed that Lantadene B, isolated from the methanolic extract of the plant, is capable of inducing cytotoxicity and G1 phase cell cycle arrest in MCF-7 cells (Shamsee et al., 2019). *Lantana camara* root extract-derived root nanoparticle exhibited cytotoxicity on MDA-MB-231 cells (Ramkumar et al., 2017). El-Din et al. also reported that the methanolic extract of *L. camara* leaves could exert cytotoxic effects on MCF-7 and MDA-MB-231 breast cancer cell lines (El-Din et al., 2022). However, the detailed mechanism of cytotoxicity on MDA-MB-231 cells was not explored. In our previous study, it was reported that the ethanolic leaf extract of *Lantana camara* is cytotoxic to both MCF-7 and MDA-MB-231 cell lines (Pal et al., 2024). Moreover, the extract could induce the G0/G1 phase cell cycle arrest, cell death, and decreased migration potential of MDA-MB-231 cells (Pal et al., 2024). These findings make *Lantana camara* leaf extract a potential candidate for breast cancer therapy and call for further exploration of the anti-tumor effect of the extract.

Plant-derived compounds can affect various intracellular processes, like epigenetic alteration, microRNA regulation, which are vital players in cancer development and progression, including breast cancer. A study by Hargraves et al. revealed that phytochemicals derived from cruciferous vegetables and sweet wormwood plant could up-regulate the expression of tumor suppressor miR34a, resulting in down-regulation of target gene CDK4 in MCF-7 and T47D cell lines (Hargraves et al., 2016). Polyphenol-Enriched Blueberry Preparation has also been reported to downregulate miR-210 in MDA-MB-231 and 4T1 cells and increase miR145 expression (Mallet et al., 2021). In an elaborate review by Ahmed et al., it has been mentioned that various plant-derived compounds like EGCG from tea, DIM from cruciferous can modulate miR-16, miR-21, and miR-92a. Other plant-derived compounds, Glyciollins, increased the expression of miR-181c and miR-181d, and many other microRNAs, and have a tumor suppressive role (Ahmed et al., 2020). Quercetin, a plant-derived flavonoid, is also known to regulate various miRNAs involved in apoptosis and cancer metastasis. A few of them include miR-16, 34a, 26b, miR-146a/b, 194, etc(Ahmed et al., 2020). Therefore, it would be no surprise for *Lantana camara* leaf extract to show a similar kind of epigenetic regulation. However, there are no reports available to date showing *Lantana camara*-mediated epigenetic regulation through miRNA in cancer.

Due to these reasons, in our current study, we were interested in exploring the effect of *Lantana camara* leaf extract on microRNA regulation in MCF-7 and MDA-MB-231 cells. For the study, miR-181a-5p was selected. miR-181a-5p is a member of the miR181 family, which is evolutionary conserved, indicating its importance (Yang et al., 2017). This miRNA is known for playing a dual role in breast cancer progression and regulation (Yang et al., 2017). Various reports have also shown the dual role of miR-181a-5p in breast cancer. Liu et al. have shown that miR-181a can induce the proliferation and invasion of breast cancer cells by targeting KLF6 and KLF15, respectively (Liu et al., 2020). Another report by Zhang et al. demonstrated that the upregulation of miR-181a-5p under hypoxia reduces epirubicin sensitivity in breast cancer cells by inhibiting EPDR1/TRPC1, thereby activating the PI3K/AKT signalling pathway (Y. Zhang et al., 2024). On the other hand, Lin et al. have shown that overexpression of miR-181a in HS578T breast cancer cells can reduce their proliferation, migration, and also induce autophagy (Lin et al., 2021). miR-181a is also predicted to target *BCL-2*, *MCL-1*, and *CDKN3* mRNAs, which are vital genes for cell survival and proliferation. The interaction of miR181a with *BCL-2* and *MCL-1* is already well established in the literature (Ouyang et al., 2012). However, no report is available on the interaction between *CDKN3* and miR-181a. Similarly, the role of *Lantana camara* in the regulation of miR181a and its combined effect on breast cancer cells is also not explored. Hence, this study may reveal a new avenue to breast cancer therapy mediated through miR-181a and *Lantana camara* extract.

## 2. Materials and methods

### 2.1. Cell culture

The MCF-7 and MDA-MB-231 cell line was purchased from the National Centre for Cell Science (NCCS), Pune, India. All cell lines were maintained in glucose-rich Dulbecco’s Modified Eagle Medium (DMEM) (HiMedia, India), which is supplemented with 10% (v/v) Fetal Bovine Serum (FBS) (HiMedia, India), Penicillin and Streptomycin antibiotic solution (HiMedia, India). Cells were grown at 37°C in a humidified incubator with 5% CO_2_. 1X Trypsin-EDTA solution (HiMedia, India) was used for cell harvesting during sub-culturing and experiments.

### 2.2. Collection of *Lantana camara* leaves and ethanolic leaf extract preparation

Healthy and fresh leaves of *Lantana camara* were collected from the Indian Institute of Science Education and Research Kolkata (latitude-22°57′50″N and longitude-88°31′28″E) campus and surroundings. The leaves of the plant were identified and authenticated by an expert botanist of the Central National Herbarium, Botanical Survey of India, West Bengal, India. A voucher specimen was deposited at their herbarium with a voucher/reference number IISER/AP/01. A detailed protocol for plant extract preparation is described and published in our last published paper on MDA-MB-231 (Pal et al., 2024). Briefly, fresh and healthy leaf of the *Lantana camara* plant is collected, washed, and shade-dried. After that, dried leaves are crushed to a fine powder and mixed with 100% molecular grade ethanol (Merck, Germany) in 1:10 ratio overnight in an orbital shaker. The mixture is filtered using filter paper and dried using a rotary evaporator, and the dry powder was reconstituted using 100% molecular grade ethanol (Merck, Germany) at the desired concentration and filtered using a syringe filter (0.2 Micron) and used for experiments. As the extract has been prepared in 100% ethanol, 0.8% (v/v) ethanol has been used as a solvent control in all the experiments.

### 2.3. Selection of treatment doses for the study

For treating breast cancer cells with the extract, different doses of the extract were selected based on the IC_50_ values of the extract for MCF-7 and MDA-MB-231 cells, already reported in our previous publication (Pal et al., 2024). As the IC_50_ value of the extract for MCF-7 (IC_50_: 91.66 µg/mL) was lower than MDA-MB-231 cells (IC_50_: 111.33 µg/mL), low treatment doses of the extract have been chosen for MCF-7 cells (5 µg/mL, 10 µg/mL, 20 µg/mL, 40 µg/mL, 80 µg/mL, 120 µg/mL) compared to MDA-MB-231 (40 µg/mL, 60 µg/mL, 80 µg/mL, 120 µg/mL,180 µg/mL) cells. However, the 24h treatment time is constant for both cell lines. As the treatment time is fixed, for observing cell cycle and cell migration, low doses (below IC_50)_ are selected. However, for the cell death assay, higher doses close to IC_50_ and above have been selected for a better understanding of the extract’s effect.

### 2.4. Bioinformatics prediction

Online available bioinformatics target prediction tools, Targetscan 8.0 (https://www.targetscan.org/mmu_80/) and miRDB (https://mirdb.org/) were used to predict the target of miR-181a-5p and to find the binding region of miR-181a-5p in the target gene.

### 2.5. Isolation of RNA and microRNA and reverse transcription quantitative real-time polymerase chain reaction (RT-qPCR)

MCF-7 and MDA-MB-231 cells were seeded in 60mm cell culture dishes at a seeding density of 0.4 × 10^6^ cells in each dish. At 60% -70% confluency, cells were treated with either 0.8% (v/v) ethanol or different doses of *Lantana camara* extract (20 µg/mL, 40 µg/mL, 60 µg/mL, 80 µg/mL, 120 µg/mL, 180 µg/mL) for 24 hours. Total RNA was extracted using total RNA extraction reagent (Takara, Japan) following the manufacturer’s protocol, which is explained in detail in our previous publication (Pal et al., 2024). For the isolation of microRNA, an additional overnight precipitation step was added using 3M sodium acetate (SRL, India) and 100% molecular grade ethanol (Merck, Germany) at -20°C temperature. The RNA sample was then treated with RNase-free DNase I enzyme (Thermo Fisher Scientific, USA) according to the manufacturer’s instructions to eliminate any residual chromosomal DNA. The samples were subsequently quantified using a microplate spectrophotometer (BioTek Epoch Microplate Spectrophotometer, Agilent Technologies Inc., USA). Total RNA was reverse transcribed into cDNA using Primescript 1^st^ strand cDNA synthesis kit (Takara, Japan) in accordance with the manufacturer’s instructions.

### 2.6. Transfection of MCF-7 and MDA-MB-231 cell lines

MCF-7 and MDA-MB-231 cells were seeded in 35mm dishes at a density of 0.2 × 10^6^ cells in each dish. At a confluency of 60% - 70%, cells were transfected with either pcDNA3.1+ or pcmiR181a using the calcium phosphate transfection method. Briefly, in a tube, 75 µL tris-EDTA buffer (pH 8) was taken. To this, 20 µL 2M CaCl_2_ (SRL, India) and 15 µg purified plasmid DNA were added. To this mixture, 110 µL 2X HBS was added dropwise under gentle vortex conditions. The mixture was allowed to stand at room temperature for 20 minutes. In the meantime, the media of the culture dishes was replaced with antibiotic-free media with 10% FBS. After that, the transfection mixture was added dropwise to the culture dishes. After 6 hours, the media was replaced with fresh complete DMEM media, and cells were harvested after 48 hours post-transfection for experiments.

### 2.7. Cell cycle phase distribution assay

MCF-7 and MDA-MB-231 cells were seeded in 35 mm cell culture dishes (0.2 × 10^6^ cells/dish) and incubated overnight. Cells were then transfected with either pcDNA 3.1+ empty vector or pcmiR181a. After 24 hours post transfection cells were treated with 20 µg/mL, 40 µg/mL, 80 µg/mL, 120 μg/mL doses of the leaf extract for MCF-7 cells and 40 μg/mL, 60 μg/mL, 120 μg/mL, 180 μg/mL doses of the leaf extract for MDA-MB-231 cells along with vehicle treated and untreated control cells. Following 24 hours of treatment, cells were harvested from the culture dishes using 1X Trypsin-EDTA and collected along with the floating cells derived from spent media. Cell pellets were washed with ice-cold 1X phosphate buffer saline (PBS). Then the cells were fixed with 70% ethanol and incubated at 4°C overnight. The next day, cells were pelleted down and washed with ice-cold 1X PBS, and were resuspended in 500 µL propidium iodide (PI) solution and incubated at 37°C for 20 minutes in the dark. Thereafter, the PI solution was removed, and cells were resuspended in 500 µL 1X PBS and analyzed using a flow cytometer (BD Biosciences, USA). Cell cycle distribution (Sub-G1, G1/G0, S, and G2/M) was determined, and the data were represented as a bar graph.

### 2.8. Detection of Cell death by flow cytometry

Double staining using the Annexin V-FITC/PI Apoptosis detection kit by BD Biosciences was performed following the manufacturer’s instructions to differentiate between apoptotic and non-apoptotic cell death. MCF-7 and MDA-MB-231 cells were seeded in 35 mm cell culture dishes (0.2 × 10^6^ cells/dish) and incubated overnight, and then were transfected with either pcDNA 3.1+ empty vector or pcmiR181a. After 24 hours post-transfection, cells were exposed to 40 μg/mL, 80 μg/mL, and 120 μg/mL doses of the leaf extract for MCF-7 cells and 60 μg/mL, 120 μg/mL, and 180 μg/mL doses for MDA-MB-231 cells, along with vehicle-treated and untreated cells, for 24 hours. Following treatment, cells were harvested along with the floating cells derived from spent media. Cells were stained following the protocol described in our previous publication (Pal et al., 2024). Briefly, the cell pellet was washed with chilled 1X PBS, followed by resuspending the cells in binding buffer. Thereafter, cells were stained with PI and Annexin V-FITC simultaneously for 20 minutes in the dark and were analyzed within an hour by a flow cytometer to obtain the scatter plot with viable cells (bottom left quadrant), early apoptotic cells (bottom right quadrant), late apoptotic cells (upper right quadrant), and non-apoptotic cells (upper left quadrant). Data was represented graphically as Bar plots.

### 2.9. Wound healing assay

MCF-7 and MDA-MB-231 cells (0.2 × 10^6^ cells/dish) were seeded in 35mm dishes and allowed to grow until a uniform monolayer of cells was formed, and then were transfected with either pcDNA 3.1+ empty vector or pcmiR181a. After 24 hours post-transfection, a wound was created in each dish using a sterile P10 microtip, and the spent media was removed from the culture dish. The monolayer of cells was washed twice with 1X PBS, and complete media was added to the cells, followed by the addition of cycloheximide (0.5µg/mL). Immediately after the addition of cycloheximide, the wounded cells were treated with 5 µg/mL, 10 µg/mL, 20 µg/mL doses of the leaf extract for MCF-7 cells and 40 μg/mL, 60 μg/mL, 80 μg/mL doses of the leaf extract for MDA-MB-231 cells, along with vehicle-treated and untreated control cells. The cells were imaged at 0 hours and 24 hours post-treatment using an inverted microscope (Evos XL Core, Invitrogen, United States) with a camera. The images were analyzed, and wound closure was calculated by ImageJ (ImageJ software, NIH, USA) using the following formula:

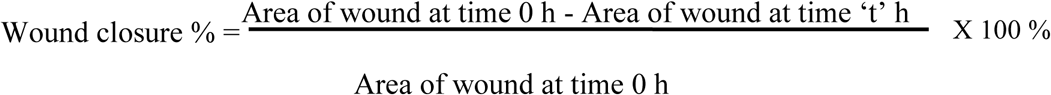

where t = specific time intervals after 0 h

### 2.10. Cloning of 3ʹ UTR of *CDKN3* in pEGFPC1 vector

A part of the *CDKN3* 3´UTR region was digested with HindIII and KpnI and run on 1% agarose gels. The digested inserts were gel-purified, quantified, and ligated to the double-digested pEGFP-C1 vector downstream *egfp* reporter gene and transformed using CaCl_2_ (100mM). All positive clones were selected from LB-agar media against kanamycin (final concentration 100 µg/mL). Now the construct is recognized as pEGFPCDKN3. The empty vectors (pcDNA3.1+, pEGFPC1) and pCmiR181a were the kind gift from Dr. Sumita Sengupta (nee Bandyopadhyay), University of Calcutta. SnapGene (www.snapgene.com) was used for the preparation of the vector map with the construct.

### 2.11. Site-directed mutagenesis of the miR-181a-5p seed region on the 3ʹ UTR of *CDKN3*

Three bases at the seed region of miR-181a-5p were altered on the 3´ UTR of the *CDKN3* mRNA using primers spanning the region (Table S1). Single primer amplification technique was used to amplify the pEGFPCDKN3 using GXL high fidelity polymerase (Takara, Japan) using the manufacturer’s protocol. After amplification, the PCR products were reannealed using a denaturation and slow cooling technique. After that, the product was purified using a DNA purification kit (Invitrogen, USA). 2µg of purified product was digested with DpnI restriction enzyme following the manufacturer’s protocol. The digested product was run on agarose gel, and the desired band was cut from the gel and purified using a gel purification kit. The purified product was used for transforming E. coli XL-1 Blue competent cells, and positive colonies were selected from LB-agar media against kanamycin (final concentration 100 µg/mL). It was further confirmed by sequencing and was named as pECDKN3SDM.

### 2.12. GFP Reporter assay

MCF-7 cells were chosen for doing reporter assay. Cells were seeded in 60mm dishes in triplicate at a density of 0.4×10^6^ cells per dish, a day before the experiment. The next day, the cells were transiently transfected by pcDNA3.1+, pEGFPC1, pcmiR181a, pEGFPCDKN3, pECDKN3SDM in antibiotic-free DMEM media (DMEM, 10%FBS), followed by changing the transfection medium after 6 hours of incubation. Cells were harvested after 48 hours using 1X trypsin-EDTA solution. Expression of GFP was observed through Western blot analysis. GAPDH was used as a loading control.

### 2.13. Western Blot analysis

MCF-7 cells were seeded in 60mm dishes at a seeding density of 0.3×10^6^ cells/dish and were grown till 60% - 70% confluency. The next day, the cells were transiently transfected by pcDNA3.1+, pEGFPC1, pcmiR181a, pEGFPCDKN3, pECDKN3SDM in antibiotic-free DMEM media (DMEM, 10%FBS), followed by changing the transfection medium after 6 hours of incubation. Cells were harvested after 48 hours using 1X trypsin-EDTA solution and were washed twice with 1X PBS. Thereafter, these cells were lysed with RIPA cell lysis buffer (Himedia, India) on ice, followed by protein estimation by the Bradford protein estimation method. 60µg of protein was run on a 12% SDS page gel at a constant 150 volts for 1.5 hours, followed by transfer on a PVDF membrane at 160mA for 1 hour. After the transfer was complete, the membrane was washed with 1X TBST and was blocked with 5% BSA in TBST for 1hour at room temperature under gentle rocking conditions. After blocking was complete membrane was again washed with 1XTBST for 5 minutes, and then incubated with primary antibody against GFP (ABclonal, AE078, 1:5000 dilution) and GAPDH (Affinity Biosciences, AF7021, 1:3000 dilution) overnight at 4°C. The next day, the membrane was again washed with 1XTBST thrice and was incubated with horseradish peroxidase conjugated secondary antibody to goat anti-rabbit IgG (1:5000, Cell Signaling Technology) for 1hour at room temperature. After that membrane was again washed with 1XTBST thrice and developed with enhanced chemiluminescence, scanned, and recorded.

### 2.14. Statistical analysis

All graphs were generated using GraphPad Prism 9. Error bars indicate mean ± SEM of at least three independent experiments. A parametric paired t-test was used for analysis of statistical significance. *P*-values <0.05 were considered to be statistically significant, while *P*>0.05 were considered non-significant (ns).

## 3. Results

### 3.1. *Lantana camara* leaf extract upregulated the expression of hsa-miR-181a-5p in MCF-7 and MDA-MB-231 cell lines

To observe the effect of *Lantana camara* treatment on the expression of miR-181a-5p in MCF-7 and MDA-MB-231 cells, miRNA was isolated and cDNA prepared after 24 hours of leaf extract treatment, and quantitative real-time PCR was done. The PCR results were represented as Bar plots (Figure 1). It is observed that both MCF-7 and MDA-MB-231 cells have a low expression of miR-181a-5p in the vehicle control condition, whereas treatment with *Lantana camara* increased the expression of miR-181a-5p in both cell lines significantly (Figure 1), with the increase in *Lantana camara* dose. For MCF-7 (Figure 1A) cells at 20 µg/mL dose, there is a non-significant increase, but at higher doses, the expression of miR-181a has increased significantly. Similarly, for MDA-MB-231 cells (Figure 1B), a significant increase in miR-181a expression was observed at 120 µg/mL and 180 µg/mL doses. From these results, it can be concluded that *Lantana camara* leaf extract could alter the expression of miR-181a in both cells, indicating a potential role of miR-181a in breast cancer.

**Figure 1.**
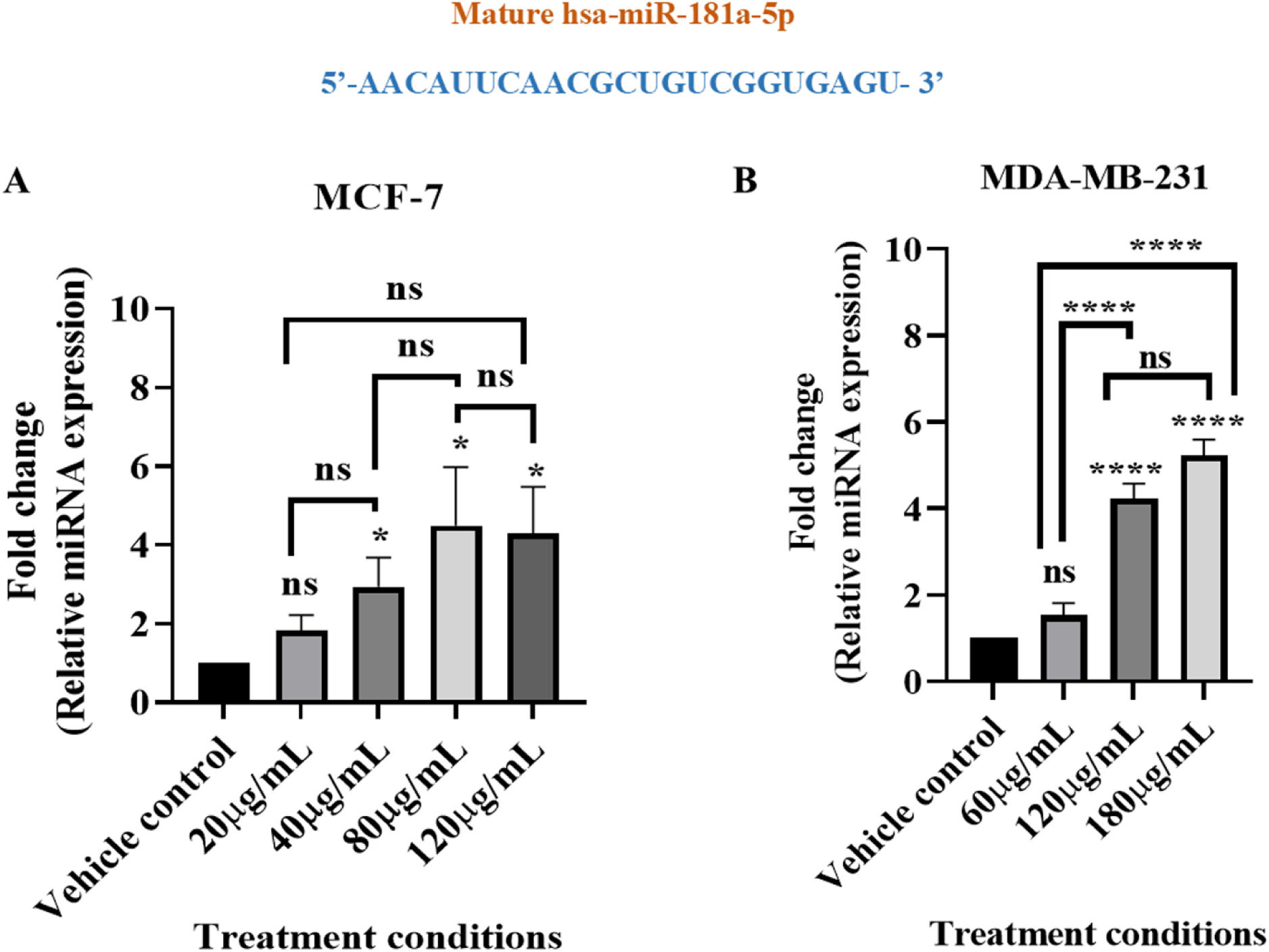
*Lantana camara* upregulated the expression of miR-181a-5p in MCF-7 and MDA-MB-231 cell lines. (A), and (B) Bar plots showing the expression of miR-181a-5p in MCF-7 and MDa-MB-231 cells. Data is plotted as mean ± SEM from three independent biological replicates. Asterisks indicate statistically significant differences when compared to vehicle control (\**P* < 0.05; \*\**P* < 0.01; \*\*\**P* < 0.001; \*\*\*\**P* < 0.0001 and ns signifies non-significant).

### 3.2. *Lantana camara* leaf extract treatment and ectopic expression of miR-181a-5p altered the mRNA level expression of miR-181a-5p target mRNAs: *BCL-2*, *MCL-1*, and *CDKN3* in MCF-7 and MDA-MB-231 cell lines

It is known that miRNA can bind to different target mRNAs and control their function by various mechanisms. To further investigate the effect of leaf extract on the hsa-miR-181a-5p targets, a few genes involved in cancer cell survival and proliferation were chosen, and changes in their mRNA level expression upon 24h *L. camara* treatment were observed through real-time quantitative PCR (Figures 2–4). Data revealed that the mRNA level expression of miR-181a-5p targets-*BCL-2*, *MCL-1*, and *CDKN3* decreased with the leaf extract treatment in both cell lines. Furthermore, to investigate the effect of ectopic expression of miR-181a on the mRNA level expression of these genes, both MCF-7 and MDA-MB-231 cells were transfected with pcDNA3.1+ and pcmiR181a plasmids, and the mRNA level expression of target genes was observed. Results revealed that the expression of *BCL-2*, *MCL-1*, and *CDKN3* mRNAs (Figures 2–4) was significantly reduced in miR-181a overexpressing MCF-7 and MDA-MB-231 cells when compared to control cells. Therefore, these results also indicate the involvement of miR181a in the regulation of these mRNAs. As the above-mentioned genes are critical players in the proliferation, survival, and migration of cells, the effects of overexpression of miR-181a-5p in the regulation of these vital parameters were also explored.

**Figure 2.**
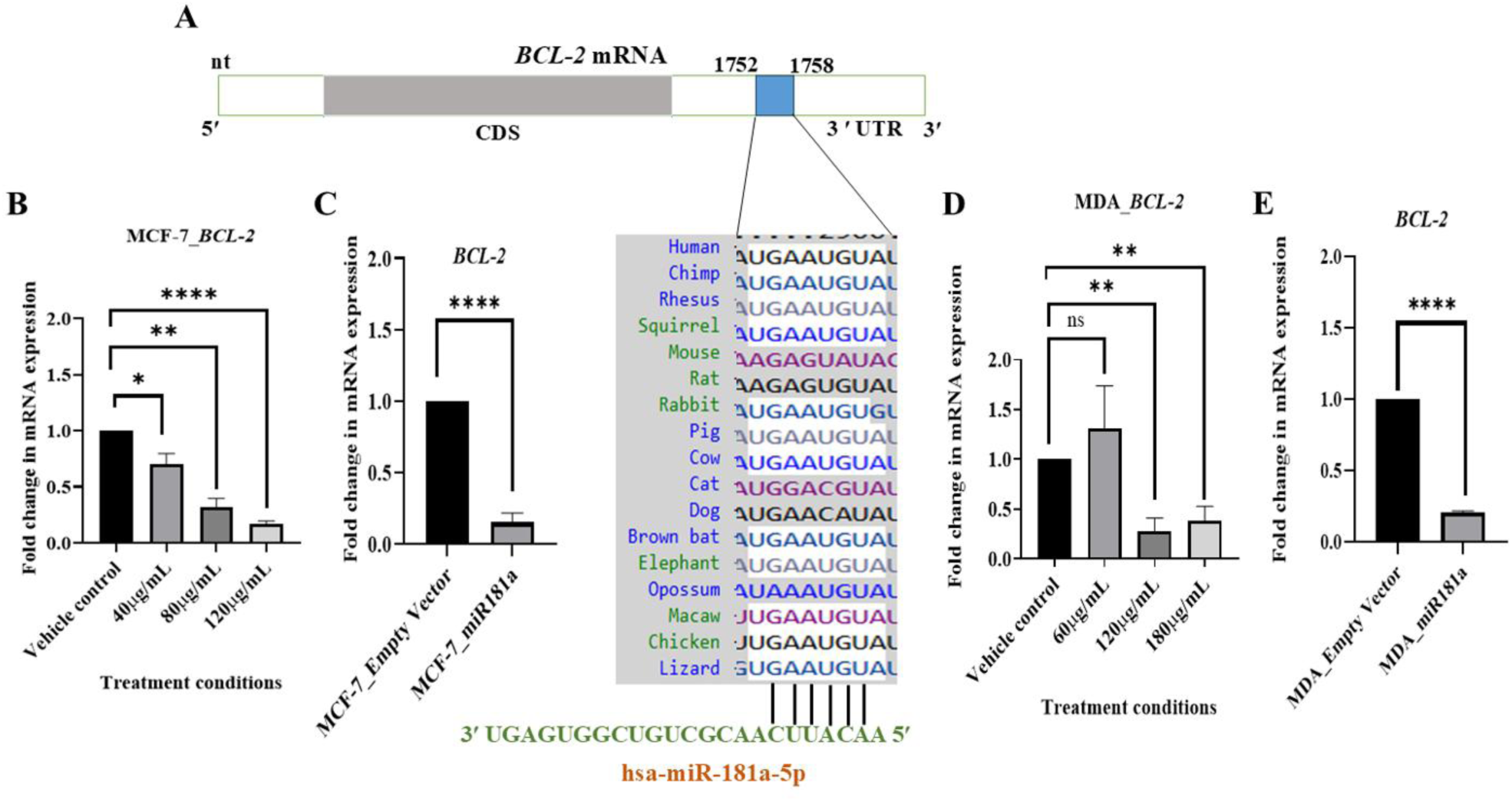
mRNA level expression of *BCL-2* decreased upon *Lantana camara* treatment and miR-181a-5p overexpression conditions in MCF-7 and MDA-MB-231 cells. (A) Schematic representation of in silico search of miR-181a-5p seed region in 3ʹ UTR of *BCL-2* mRNA and its cross-species conservation. (B), and (D) bar plots showing the mRNA level expression of *BCL-2* in MCF-7 and MDA-MB-231 under *Lantana camara* treatment conditions. (C) and (E) bar plots showing the mRNA level expression of *BCL-2* in MCF-7 and MDA-MB-231 under miR-181a-5p overexpression conditions. Data is plotted as mean ± SEM from three independent biological replicates. Asterisks indicate statistically significant differences when compared to control (\**P* < 0.05; \*\**P* < 0.01; \*\*\**P* < 0.001; \*\*\*\**P* < 0.0001 and ns signifies non-significant).

**Figure 3:**
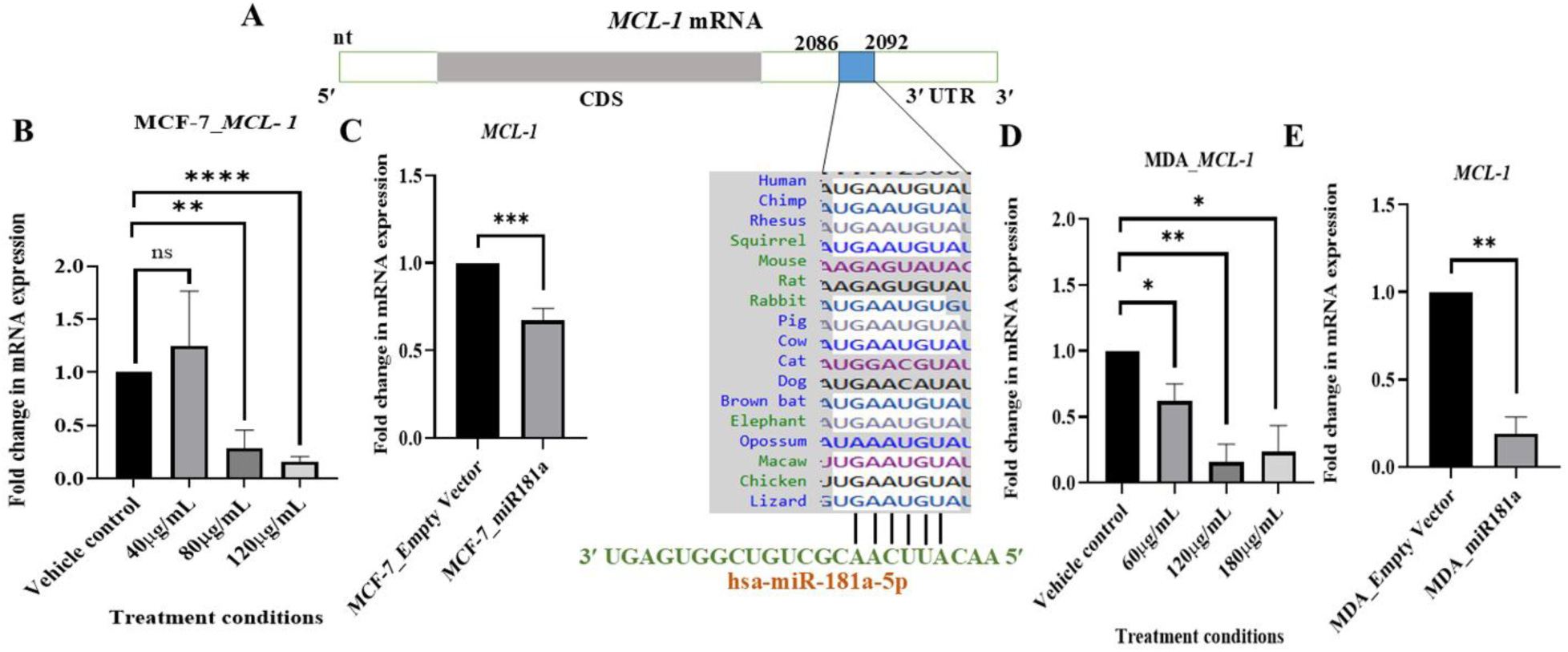
mRNA level expression of *MCL-1* decreased upon *Lantana camara* treatment and miR-181a-5p overexpression conditions in MCF-7 and MDA-MB-231 cells. (A) Schematic representation of in silico search of miR-181a-5p seed region in 3ʹ UTR of *MCL-1* mRNA and its cross-species conservation. (B), and (D) bar plots showing the mRNA level expression of *MCL-1* in MCF-7 and MDA-MB-231 under *Lantana camara* treatment conditions. (C), and (E) bar plots showing the mRNA level expression of *MCL-1* in MCF-7 and MDA-MB-231 under miR-181a-5p overexpression conditions. Data is plotted as mean ± SEM from three independent biological replicates. Asterisks indicate statistically significant differences when compared to control (\**P* < 0.05; \*\**P* < 0.01; \*\*\**P* < 0.001; \*\*\*\**P* < 0.0001 and ns signifies non-significant).

**Figure 4.**
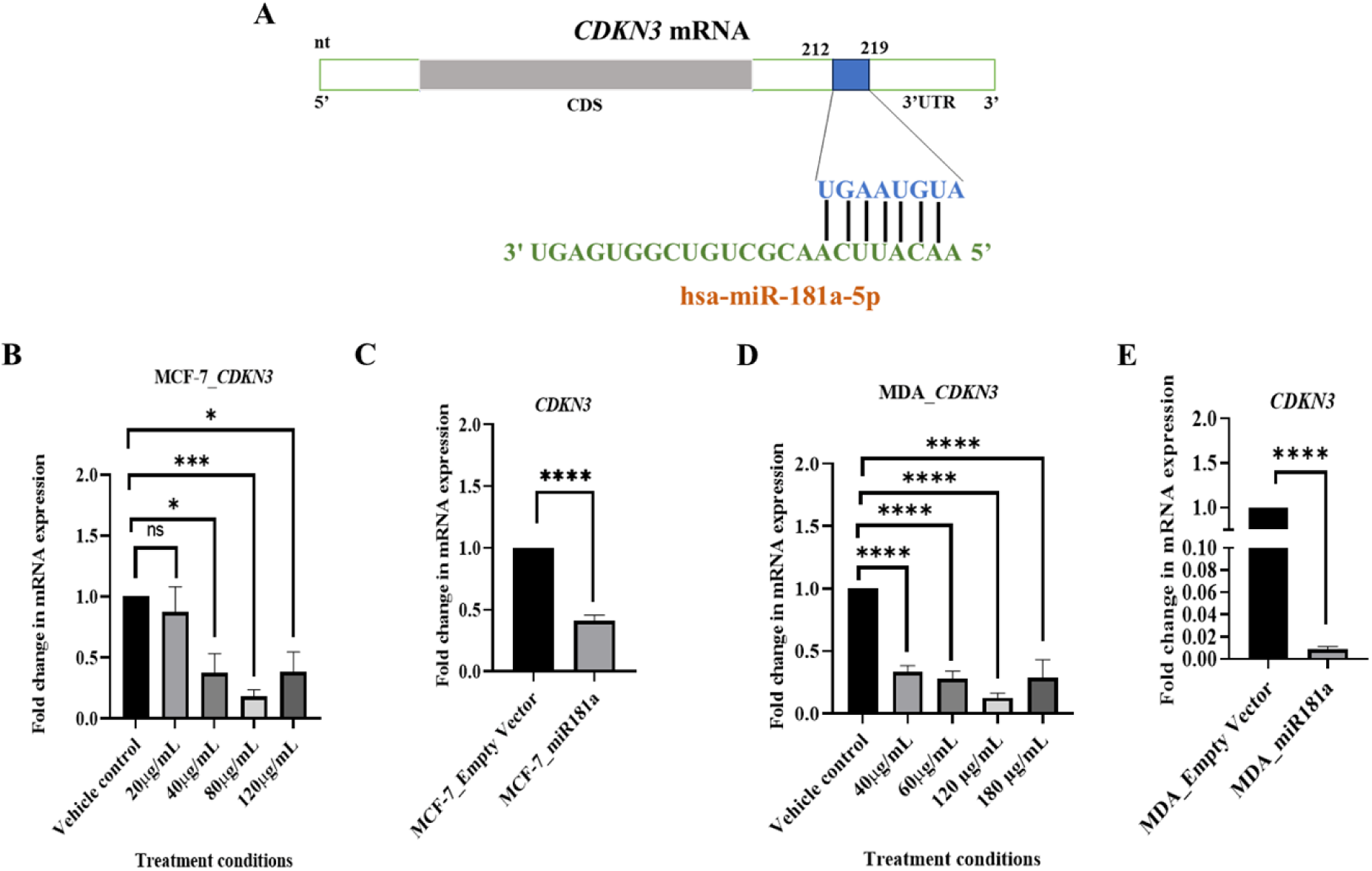
mRNA level expression of *CDKN3* decreased upon *Lantana camara* treatment and miR-181a-5p overexpression conditions in MCF-7 and MDA-MB-231 cells. (A) Schematic representation of in silico search of miR-181a-5p seed region in 3ʹ UTR of *CDKN3* mRNA. (B), and (D) bar plots showing the mRNA level expression of *CDKN3* in MCF-7 and MDA-MB-231 under *Lantana camara* treatment conditions. (C), and (E) bar plots showing the mRNA level expression of *CDKN3* in MCF-7 and MDA-MB-231 under miR-181a-5p overexpression conditions. Data is plotted as mean ± SEM from three independent biological replicates. Asterisks indicate statistically significant differences when compared to control (\**P* < 0.05; \*\**P* < 0.01; \*\*\**P* < 0.001; \*\*\*\**P* < 0.0001 and ns signifies non-significant).

### 3.3. Ectopic expression of miR-181a-5p increased sub-G1 population in both MCF-7 and MDA-MB-231 cells in the presence of *Lantana camara* leaf extract

Cell cycle analysis provides insight into cell growth, proliferation, and response to various stimuli and treatments. Therefore, observation of the distribution of cells through different cell cycle phases helps to understand the effect of overexpression of miR-181a-5p in the presence and absence of the leaf extract. In our previous publication, it has already been established that *Lantana camara* extract was able to induce G0/G1 phase arrest in MDA-MB-231 cells. Here, we intend to show whether the overexpression of miR-181a-5p could further enhance the effects of the extract. Figures 5 and 6 show the representative images of the MCF-7 and MDA-MB-231 cells in different phases of the cell cycle (sub-G1, G0/G1, S-phase, and G2/M). Data has been analyzed and plotted as Bar plots (Figure 5C, 5D, and 6C, 6D). Results of the experiment showed that in both MCF-7 and MDA-MB-231 cell lines, there is an increase in the sub-G1 population in miR-181a overexpressed cells compared to control cells (MCF-7_empty vector and MDA_empty vector) (Tables S2 and S3). This sub-G1 phase population increase was more for MCF-7_miR181a than MDA_miR181a cells (Tables S2 and S3), and the presence of *Lantana camara* extract further increased the sub-G1 accumulation in both cell lines. This increase in the Sub-G1 population is indicative of DNA fragmentation and cellular death. For the MCF-7 cell line increase in the sub-G1 population was dose dependent, followed by a subsequent decrease in G0/G1, S-phase, and G2/M phase of the cell cycle (Table S2). For the MDA-MB-231 cell line, only at the 180 µg/mL dose, there is a significant increase in sub-G1 population, indicating the highest DNA fragmentation at this condition, when miR-181a was transiently expressed compared to the control of the MDA_pcmiR181a group, with non-significant changes in other phases of the cell cycle. However, the increase in the sub-G1 population was higher in all the conditions in MDA_miR181a cells than in the MDA_empty vector cells (Table S3). An increase in the sub-G1 population in both cell lines gives a clear hint of the cell death in both cell lines.

**Figure 5.**
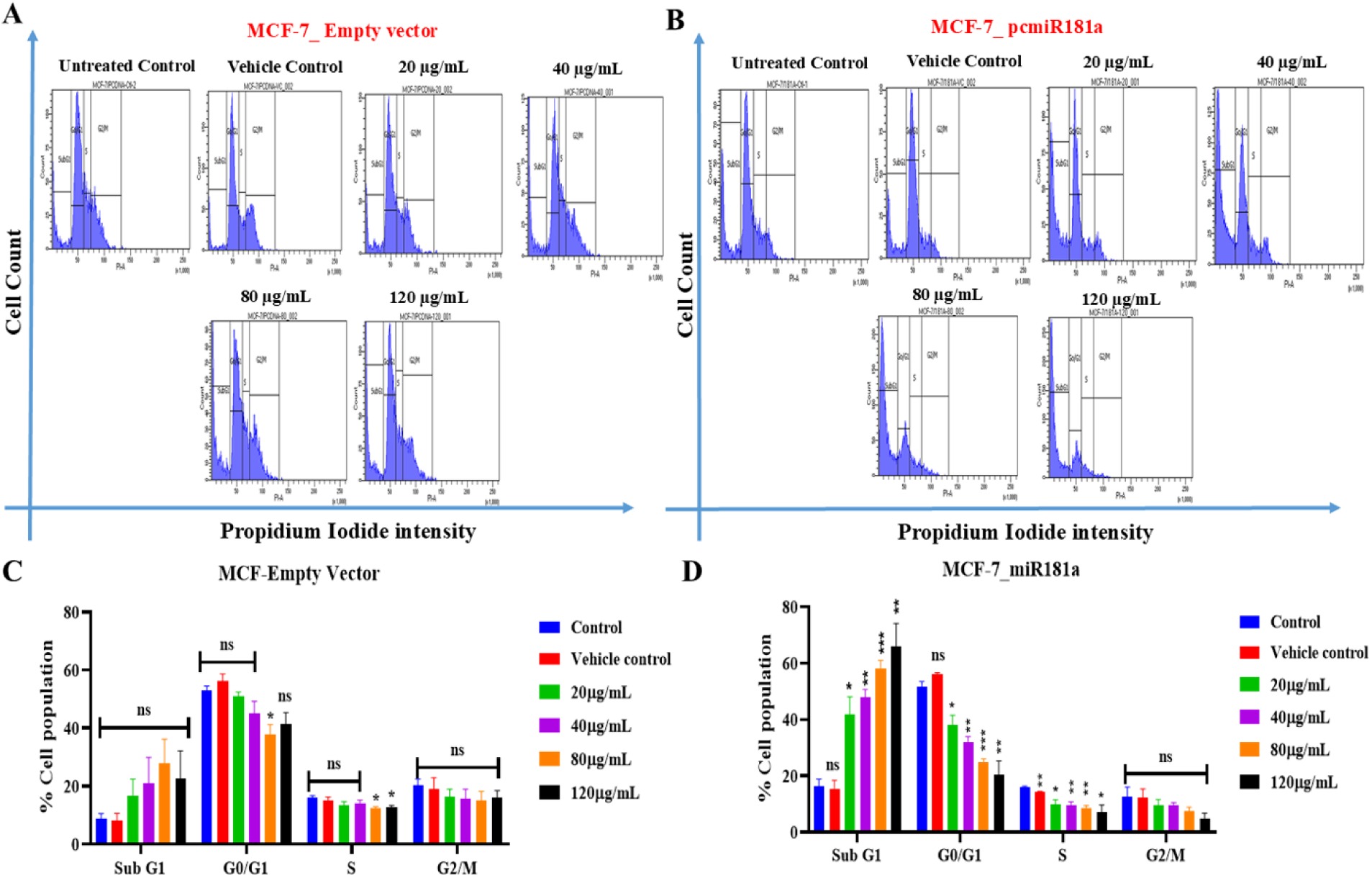
Ectopic expression of miR-181a-5p in association with *Lantana camara* leaf extract increased sub-G1 population in MCF-7 cells. (A) and (B) are the representative images showing the distribution of cells in sub-G1, G0/G1, S, and G2/M phases of the cell cycle under different treatment conditions. (C) and (D) graphical representation of the data obtained from cell cycle phase distribution assay. Data is plotted as mean ± SEM from three independent biological replicates. Asterisks indicate statistically significant differences when compared to control (\**P* < 0.05; \*\**P* < 0.01; \*\*\**P* < 0.001 and ns signifies non-significant).

**Figure 6.**
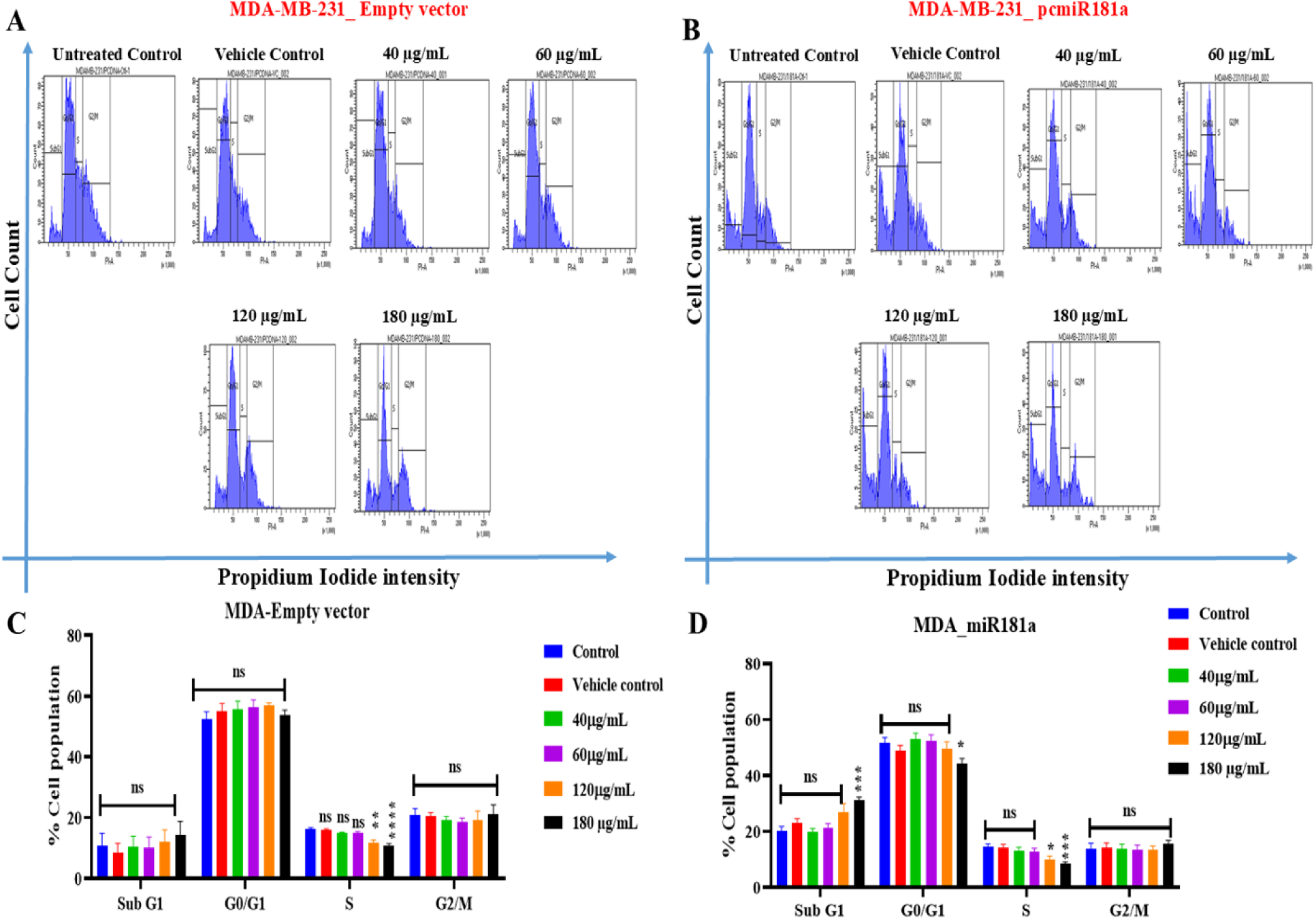
Ectopic expression of miR-181a-5p in association with *Lantana camara* leaf extract increased sub-G1 population in MDA-MB-231cells. (A) and (B) are the representative images showing distribution of cells in sub-G1, G0/G1, S and G2/M phases of cell cycle under different treatment conditions. (C) and (D) graphical representation of the data obtained from cell cycle phase distribution assay. Data is plotted as mean ± SEM from three independent biological replicates. Asterisks indicate statistically significant differences when compared to control (\**P* < 0.05; \*\**P* < 0.01; \*\*\**P* < 0.001; \*\*\*\**P* < 0.0001 and ns signifies non-significant).

### 3.4. Ectopic expression of miR-181a-5p induced cell death in MCF-7 and MDA-MB-231 cells in the presence and absence of *Lantana camara* leaf extract

An increase in the sub-G1 population is indicative of DNA breakage and hints towards cell death. To further confirm the cell death in both MCF-7 and MDA-MB-231 cells and to observe the mode of cell death, Annexin V-FITC/PI double staining was performed. Figures 7 and 8 represent the results of the experiment summarized in Table S4 and S5. For MCF-7 cells, it was observed that MCF-7 cells with transient overexpression of miR-181a-5p showed a significant increase in cell death in untreated and vehicle-treated conditions when compared to MCF-7 cells transfected with an empty vector. In untreated control condition for MCF-7_pcmiR181a cells % death cell was observed to be 5.01 ± 0.54% (early apoptotic), 19.23 ± 3.35 % (late apoptotic), 13.93 ± 0.99 (non-apoptotic), whereas for MCF-7_Empty vector, it was found to be 2.28 ± 0.40 % (early apoptotic), 6.33 ± 2.53 % (late apoptotic), 7.81 ± 1.67 % (non-apoptotic) cell death which was significantly low compared to MCF-7_pcmiR181a. The death cell population increased further with the treatment of *L. camara* extract in the MCF-7_pcmiR181a cells when compared to MCF-7_empty vector. However, the increase in cell death was not dose-dependent. In MCF-7_empty vector cells, there was a dose-dependent increase in cell death. Further comparison between MCF-7_empty vector and MCF-7_pcmiR181a (Table S4) revealed a significant increase in cell death at 120 µg/mL dose in the late apoptotic cell death quadrant (20.83 ± 1.41 % to 28.88 ± 2.43 %).

**Figure 7.**
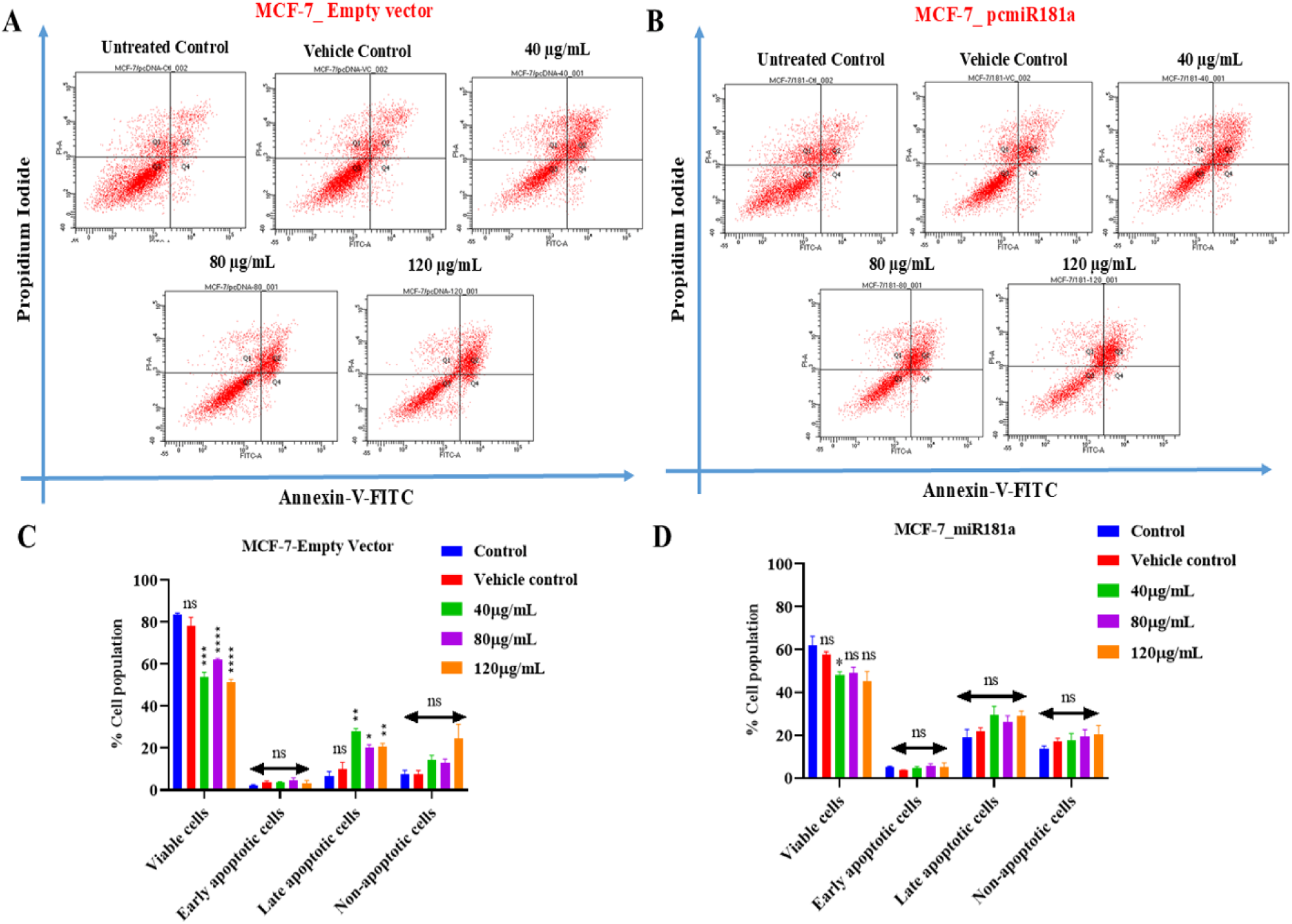
Ectopic expression of miR-181a-5p in association with *Lantana camara* leaf extract induced cell death in MCF-7 cells. (A) and (B) are the representative scatter plots showing distribution of viable (bottom left quadrant), early apoptotic (bottom right quadrant), late apoptotic (upper right quadrant), and non-apoptotic (upper left quadrant) cell populations under different treatment conditions. (C) and (D) graphical representation of the data obtained from cell death assay. Data is plotted as mean ± SEM from three independent biological replicates. Asterisks indicate statistically significant differences when compared to control (\**P* < 0.05; \*\**P* < 0.01 and ns signifies non-significant).

**Figure 8.**
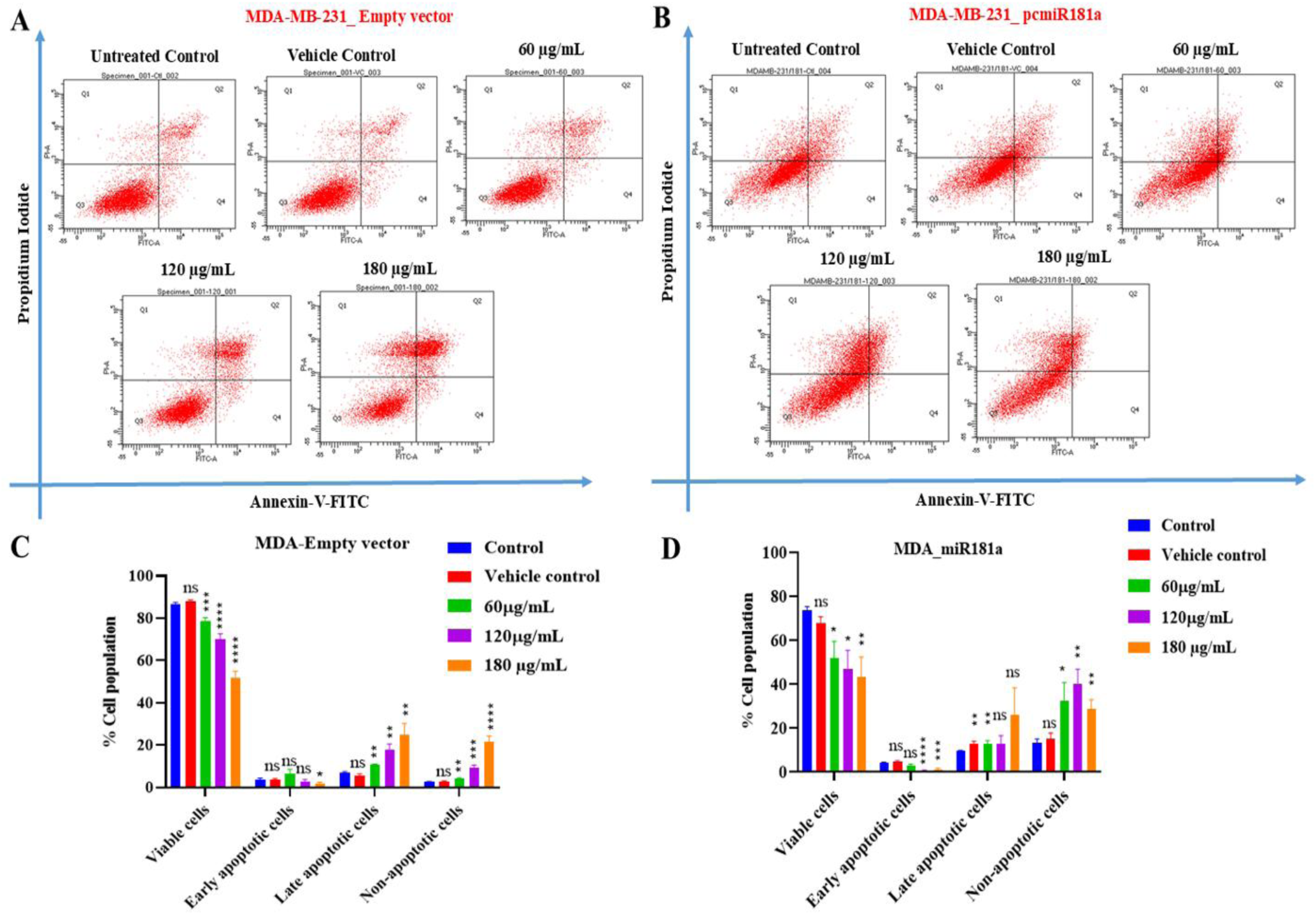
Ectopic expression of miR-181a-5p in association with *Lantana camara* leaf extract induced cell death in MDA-MB-231 cells. (A) and (B) are the representative scatter plots showing distribution of viable (bottom left quadrant), early apoptotic (bottom right quadrant), late apoptotic (upper right quadrant), and non-apoptotic (upper left quadrant) cell populations under different treatment conditions. (C) and (D) graphical representation of the data obtained from cell death assay. Data is plotted as mean ± SEM from three independent biological replicates. Asterisks indicate statistically significant differences when compared to control (\**P* < 0.05; \*\**P* < 0.01 and ns signifies non-significant).

For the MDA-MB-231 cell line (Figure 8A, 8B), it was observed that transient expression of miR-181a-5p alone was able to induce apoptotic cell death (early and late apoptotic) in untreated control and vehicle control conditions when compared to MDA_empty vector cells (Table S5). Interestingly, the introduction of *L. camara* extract had shifted the death cell population significantly towards non-apoptotic cell death, indicating that the combined effect of miR-181a-5p and *Lantana camara* plant extract could also trigger other types of cell death pathways besides apoptosis. In conclusion, miR-181a-5p could induce both apoptotic and non-apoptotic cell death significantly in both MCF-7 and MDA-MB-231 cell lines in the presence and absence of the leaf ethanolic extract.

### 3.5. Ectopic expression of miR-181a-5p inhibited cell migration in MCF-7 and MDA-MB-231 cells in the presence of *Lantana camara* leaf extract

One of the characteristics of cancer is its ability to migrate to distant parts of the body from the original tumor site. Breast tumor, specifically the triple-negative subtype, is known for its migratory ability. Breast cancer can often migrate to the bones, lungs, kidneys, and brain (Ibragimova et al., 2023). MDA-MB-231, being a triple-negative breast cancer cell, has good migratory potential. MCF-7 cells are luminal A-type breast cancer cells and do not have good migration ability, but can still migrate to nearby bones and lymph nodes. Hence, studying the migration of cancer cells is of great importance for the development of effective therapy. To study the effect of miR-181a-5p on MCF-7 and MDA-MB-231 cells, a scratch assay was performed after transiently overexpressing miR-181a-5p in both cell lines. The results of the experiment have been summarized in Figures 9 and 10. For the MCF-7 cell line (Figure 9), it has been observed that the migration of the cells has reduced significantly in both MCF-7_Empty vector and MCF-7_miR181a conditions with the increase in the *Lantana camara* dose, and the effect was maximum at 20 µg/mL dose. Comparing MCF-7_Empty vector and MCF-7_miR181a conditions, it was also observed that miR181a could not reduce the migration of the cells alone, but the introduction of the leaf extract reduced migration significantly (Figure 9C). In MDA-MB-231 cells, similar effects were observed with maximum inhibition of migration at 80 µg/mL dose (Figure 10B, 10C). From the above findings, it can be concluded that overexpression of miR-181a alone could not inhibit the migration of the breast cancer cells efficiently, but in the presence of the *L. camara* leaf extract, miR-181a reduced the migration potential of breast cancer cells significantly.

**Figure 9.**
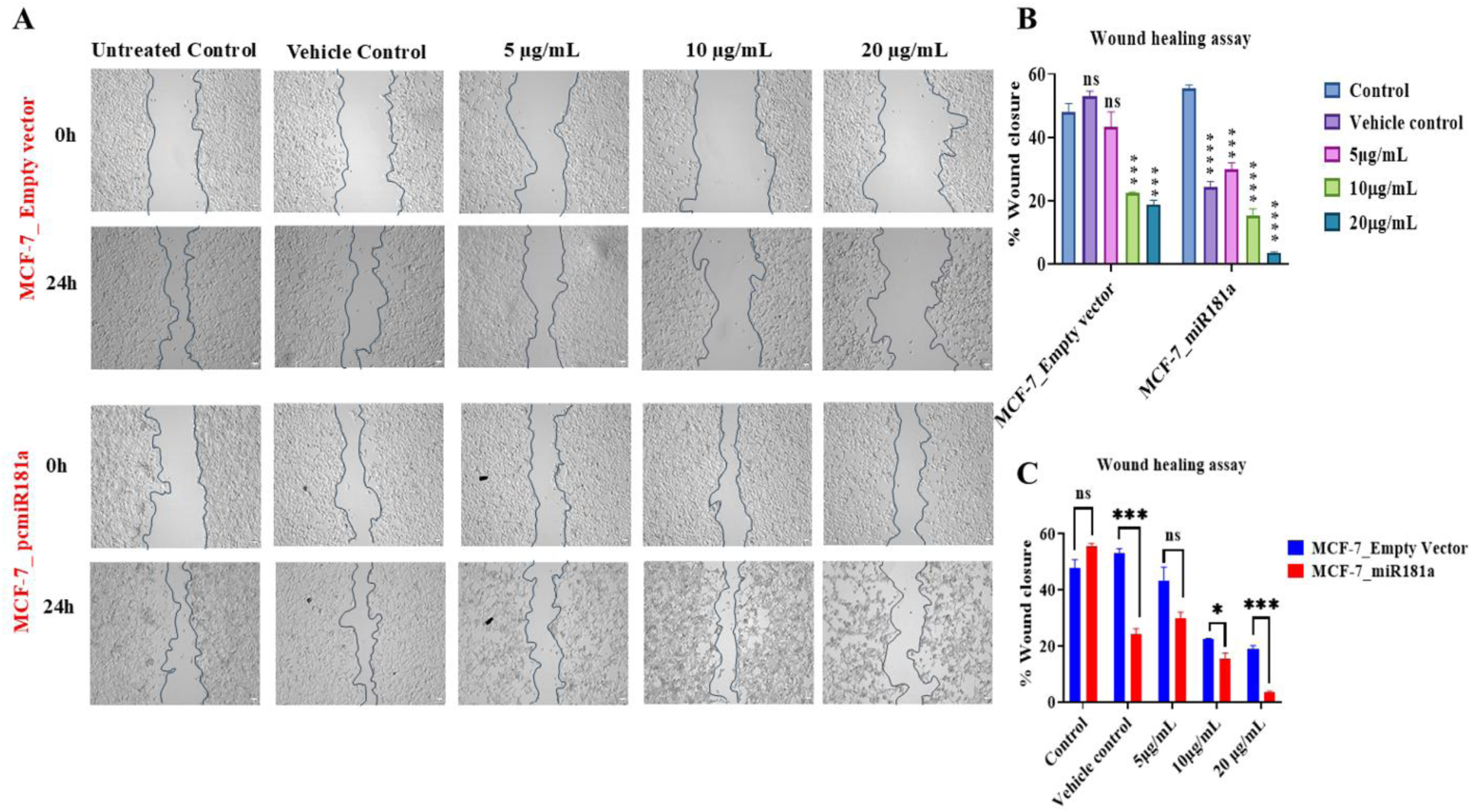
Ectopic expression of miR-181a-5p in association with *Lantana camara* leaf extract inhibited migration in MCF-7 cells. (A) For the wound healing assay, untreated, vehicle-treated and extract-treated MCF-7 cells (transfected with either empty vector (upper panel) or pcmiR181a (lower panel)) were subjected to a scratch at monolayer condition and then incubated for another 24 hours. Wound closure was visualized using a phase contrast microscope, and images were captured at 0 h and 24 h under 10X magnification. (B), (C) Bar diagram representing wound closure potential of MCF-7 cells after 24h. Data is represented as means ± SEM of three independent biological replicates, where * is *P*≤0.05, ** *P*≤0.01, *** *P*≤0.001, and ns signifies non-significant.

**Figure 10.**
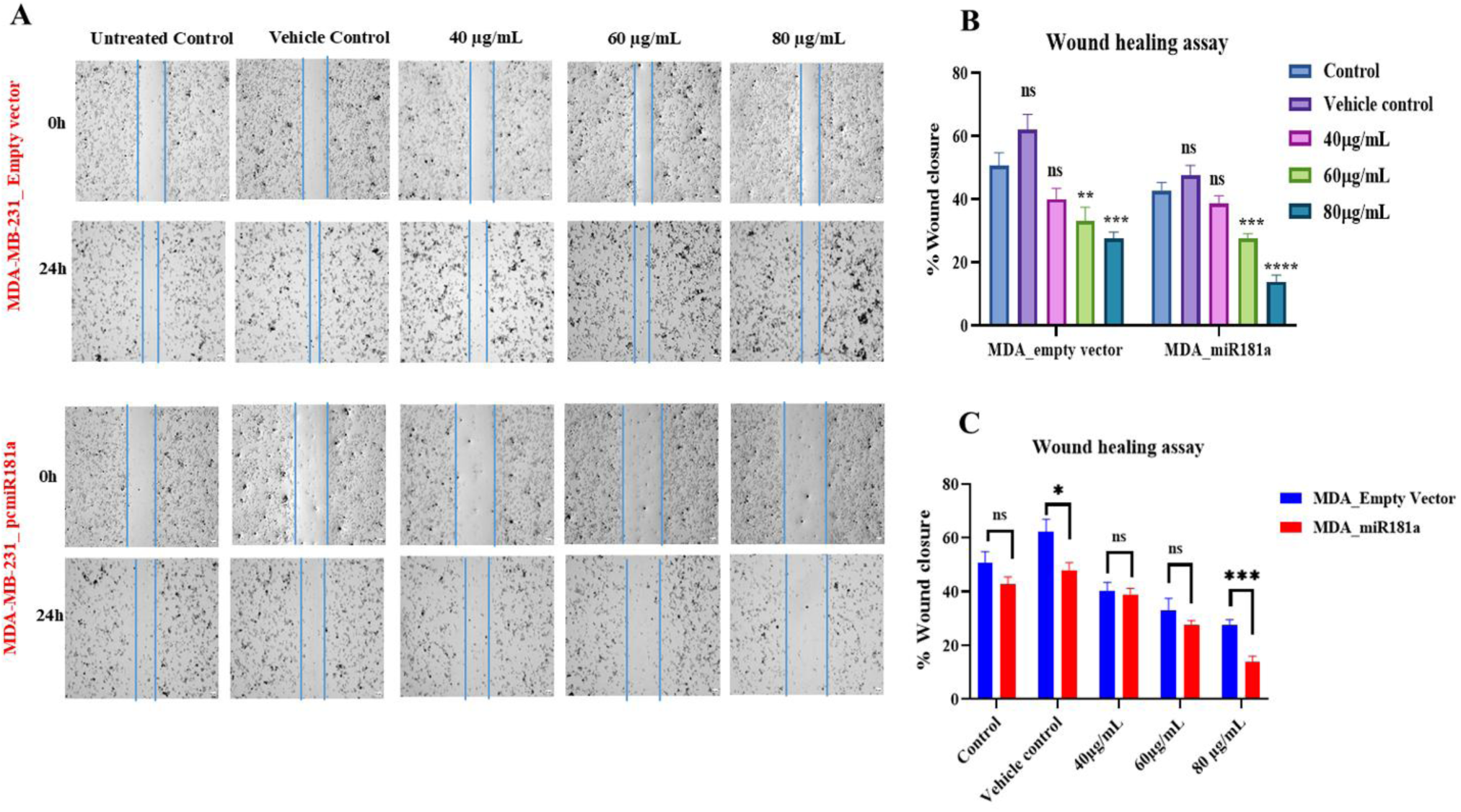
Ectopic expression of miR-181a-5p in association with *Lantana camara* leaf extract inhibited migration in MDA-MB-231 cells. (A) For the wound healing assay, untreated, vehicle-treated and extract-treated MDA-MB-231 cells (transfected with either empty vector (upper panel) or pcmiR181a (lower panel)) were subjected to a scratch at monolayer condition and then incubated for another 24 hours. Wound closure was visualized with a phase contrast microscope, and images were captured at 0h and 24h under 10X magnification. (B), (C) Bar diagram representing wound closure potential of MDA-MB-231 cells after 24h. Data is represented as means ± SEM of three independent biological replicates, where * is *P*≤0.05, ** *P*≤0.01, *** *P*≤0.001, and ns signifies non-significant.

### 3.6. miR-181a-5p targets the 3ʹ untranslated region of the *CDKN3* mRNA

*In silico* target scan (Target scan Human version 8) revealed the binding site of hsa-miR-181a to the 3ʹ UTR of *CDKN3* mRNA. To confirm the interaction between hsa-miR-181a and the *CDKN3* gene, a reporter assay was performed. MCF-7 cells were transiently transfected with pEGFPC1+pcDNA3.1+, pEGFPCDKN3+pcDNA3.1+, pEGFPCDKN3+pcmiR181a, pECDKN3SDM + pcmiR181a (Figure 11). After 48h, cells were harvested, and a whole cell lysate was prepared to check the protein level expression of the GFP reporter gene in all the conditions. It was observed that under pEGFPC1+pcDNA3.1+ and pEGFPCDKN3+pcDNA3.1+ conditions, the expression of GFP was quite high, which was significantly reduced under pEGFPCDKN3+pcmiR181a conditions (Figure 11D, 11E). The expression of GFP was partially regained when the mutated *CDKN3* construct was present, i.e, pECDKN3SDM + pcmiR181a condition. Hence, it can be concluded from the findings that miR-181a-5p can specifically bind to *CDKN3* 3’ UTR and result in the translational repression of *CDKN3*. Figure 11C, Bar plots representing the relative mRNA expression of *CDKN3* in MCF-7 and MDA-MB-231 cells.

**Figure 11:**
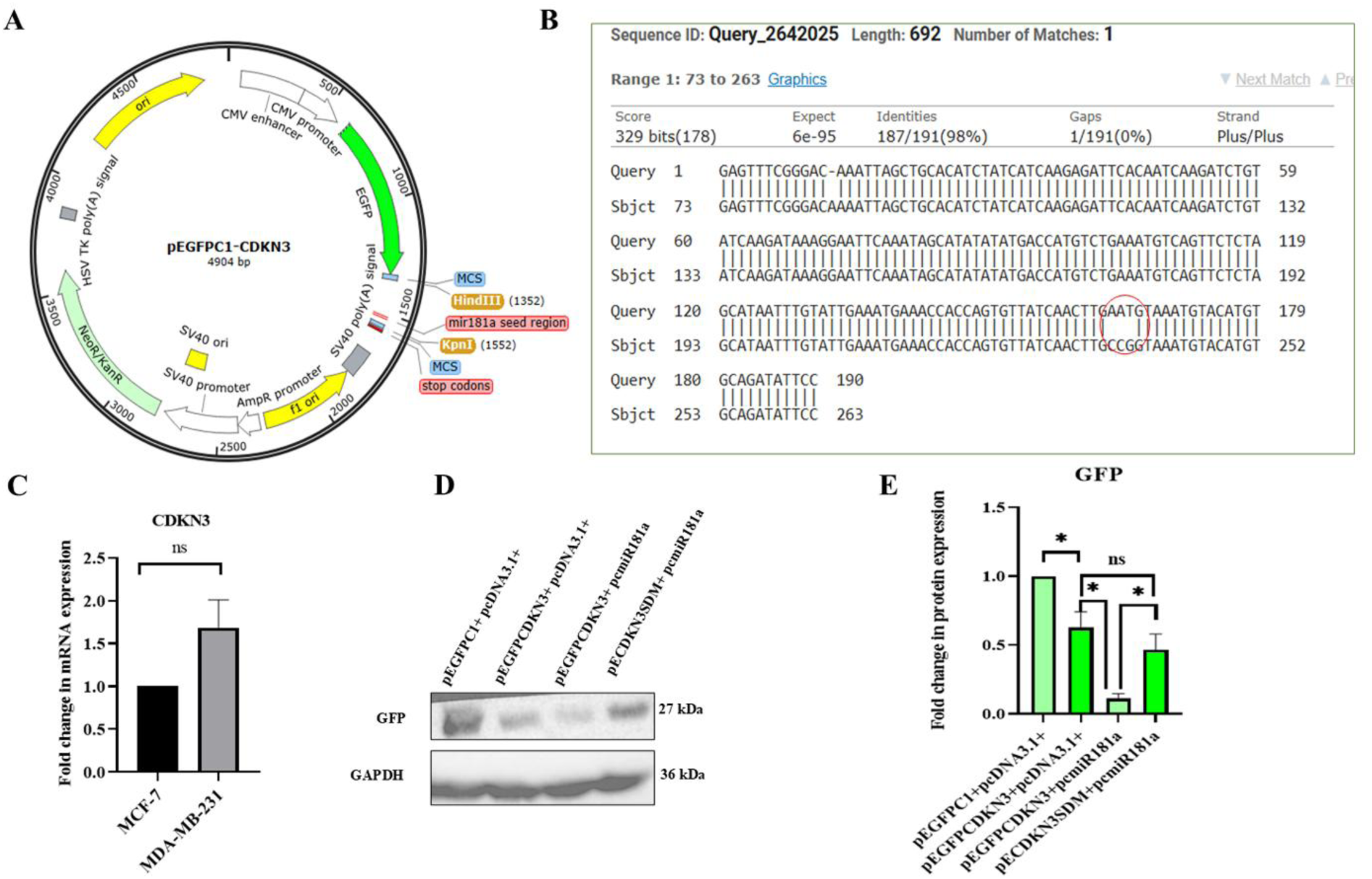
miR-181a-5p targets the 3ʹ UTR of the *CDKN3* mRNA. (A) Vector map of pEGFPCDKN3 with cloned 3ʹ UTR of *CDKN3* mRNA with miR-181a-5p seed region. (B) Alignment of wild-type *CDKN3* 3ʹ UTR sequence with mutated 3ʹ UTR in the miR-181a-5p seed region (red circle). (C) Relative expression of *CDKN3* mRNA in MCF-7 and MDA-MB-231. (D) Representative Western blot pictures for GFP and GAPDH (loading control) (E) Bar plot showing the results of Western blot. Data is plotted as mean ± SEM from three independent biological replicates. Asterisks indicate statistically significant differences when compared to the control (\**P* < 0.05, ns signifies non-significant).

## 4. Discussion

Breast cancer is one of the predominant causes of death among females globally. Among different subtypes of breast cancer, Luminal A type is the most predominant (60%) and has a better prognosis and overall survival (Burguin et al., 2021). Triple-negative breast cancer, on the other hand, is the most invasive type of breast cancer and has a worse prognosis and lower life expectancy (Burguin et al., 2021). Treatment options for breast cancer are also limited, mainly for triple-negative breast cancer, due to the absence of hormone receptors. For luminal A-type cancer, although there are various treatment options like surgery, hormone therapy, and chemotherapy but all these come with several long-term side effects (Burguin et al., 2021). For triple-negative breast cancer, chemotherapy is the only treatment option, but excessive use of chemotherapy gives rise to drug resistance and severe side effects (Mahmoud et al., 2022). To overcome these challenges, natural products isolated from plants can be an alternative option. Several breast cancer drugs, which are derived from plants like Taxol, Paclitaxel, vincristine, and sulforaphane etc, are effectively used for breast cancer treatment. Recent studies have also reported that various plant-derived compounds like genistein, resveratrol, curcumin, and lycopene have exhibited their cytotoxic and antimigratory effects on triple-negative breast cancer (Ke et al., 2022). In our preliminary work, it has been reported that *Lantana camara* leaf ethanolic extract exhibits significant cytotoxicity in MCF-7 and MDA-MB-231 cells. The extract has also induced G0/G1 phase cell cycle arrest in MDA-MB-231 cells, resulting in DNA condensation in the nucleus, followed by cell death(Pal et al., 2024). Moreover, *L. camara* can also inhibit cell migration in the aggressive MDA-MB-231 cells (Pal et al., 2024). All these findings point towards the therapeutic potential of the extract. To further elucidate the detailed underlying mechanism of the extract, we have checked the expression of a vital miRNA, miR-181a, after *L. camara* treatment. miRNAs are small RNA molecules (17-25 nucleotides) that play a vital role in breast cancer and other cancers. miRNAs have the inherent property to target different mRNA molecules and inhibit their activity either by repressing translation or by sequestering into the P-body (O’Brien et al., 2018). miR-181a-5p is a well-known microRNA that plays a pivotal role in various types of cancers, including breast cancer (Martino et al., 2025). Not only that, recently miRNAs are emerging as an effective therapeutic tool for cancer therapy. In the current study, it was found that *L. camara* upregulated the expression of miR-181a in both the MCF-7 (Figure 1) and MDA-MB-231 (Figure 1) cell lines, and also downregulated the expression of miR-181a target genes, including *BCL-2*, *MCL-1*, and *CDKN3*. *BCL-2* and *MCL-1* (Figures 2, 3) are two vital anti-apoptotic genes that play a crucial role in resisting cell death (Roberts et al., 2021). Downregulation of these genes gives a hint towards cell death, whereas the *CDKN3* (Figure 4) gene is an important regulator of cell proliferation and migration in cancer (C. Zhang et al., 2024). Overexpression of miR-181a in MCF-7 and MDA-MB-231 cell lines also downregulated the same expression of the above genes.

To delineate the role of miR-181a in the survival and migration of breast cancer cells, in combination with *Lantana camara* cell cycle phase distribution assay, Annexin V-FITC/PI apoptosis assay, and wound healing assay, was performed. From the results of experiment 3.3, it can be concluded that miR181a alone and in the presence of *L. camara* extract could induce DNA fragmentation, which is evident by the increase in the Sub-G1population in both cell lines (Table S2 and S3). This increase in Sub-G1 population is an indication of cell death, and it was further confirmed by performing Annexin V-FITC/PI double staining. Results of the experiment (Figures 7, 8) supported the findings of the previous experiment 3.3.

miR-181a alone and in the presence of the extract caused significant cell death in both cell lines (Figures 7 and 8) in both apoptotic and non-apoptotic pathways. Non-apoptotic death was more prevalent in MDA-MB-231 cells transiently expressing miR-181a under leaf extract treatment conditions. Non-apoptotic population of death cells may indicate different cell death pathways like necrosis, pyroptosis, necroptosis, ferroptosis, perthonatos, autophagy, etc (Zhang et al., 2025). Hence, Further exploration of these may give deeper insight into the cell death mechanism triggered in the MDA_pcmiR181a cells in the presence of *Lantana camara* extract. One of the features of cancer cells is their ability to metastasize to distant parts of the body from the primary tumor site, and it is more predominant in late-stage cancer. Although Luminal A subtype is less invasive in nature; these cancer cells can migrate to the bones (Fan et al., 2023). MCF-7 is luminal A type breast cancer cell line, and MDA-MB-231 is a highly invasive, late-stage triple-negative breast cancer cell line (Welsh, 2013); therefore, it was important to check the effect of miR-181a overexpression on cancer cell migration. Here, in this study, it has been observed that miR-181a was able to inhibit the migration of both MCF-7 and MDA-MB-231 cell lines in the presence of *Lantana camara* extract (Figures 9, 10). However, miR-181a could not inhibit migration alone. Hence, it can be concluded that the combined effect of miR-181a and the extract could also impede the migratory property of both cell lines. As already mentioned earlier, miRNA generally binds to the 3ʹ UTR of target mRNAs and controls their function. In the literature, it is reported that miR-181a can target the 3ʹ UTR of *BCL-2* and *MCL-1* and regulate their function (Ouyang et al., 2012). However, no report is available on the binding of miR-181a-5p to *CDKN3*. To validate that miR-181a can target *CDKN3* 3ʹ UTR, a GFP-reporter assay was done, and the result (Figure 11) confirms their interaction. In this study, we have examined three crucial parameters of cancer cells: cell proliferation, survival, and migration, and found that overexpression of miR-181a-5p was able to significantly affect all three parameters either alone or in the presence of the leaf ethanolic extract.

All these findings make *Lantana camara* leaf extract and miR181a potential candidates for breast cancer treatment in the future. However, future studies to identify the bioactive compound(s) in *Lantana camara* leaf extract and an *in vivo* study to delineate its effect may open new paths for breast cancer management.

## 5. Conclusion

To summarize, we report that *Lantana camara* upregulated the expression of miRNA-181a-5p and downregulated some important target mRNAs of the miR-181a. Transient overexpression of miR-181a in breast cancer cells also showed the similar results. Moreover, overexpression of miR-181a was able to induce sub-G1 accumulation and cell death, and it increased in the presence of the leaf extract. miR-181a overexpression also impeded cell migration in the presence of the extract in both MCF-7 and MDA-MB-231 cells. All these findings together make *Lantana camara* and miR-181a-5p potential candidates for breast cancer management in the future.

## Authors’ contributions

**Arundhaty Pal:** Conceptualisation, Formal analysis, Investigation, Methodology, Writing – original draft. **Sourav Sanyal:** Formal analysis, Investigation, Methodology, Writing – original draft. **Tapas Kumar Sengupta:** Conceptualisation, Funding acquisition, Project administration, Supervision, Writing – original draft.

## Availability of Data and Materials

In this study, all data generated or analyzed are included in the article and supplementary section.

## Ethical Approval

Not applicable.

## Consent to Participate

All authors had consented to participate in the study.

## Consent for Publication

All authors have given consent for publication.

## Acknowledgement

AP, SS, and TKS acknowledge the Indian Institute of Science Education and Research Kolkata (IISER Kolkata) for the research facilities. AP and SS acknowledge the Department of Biotechnology, Government of India (DBT-India) for the research fellowship. All authors would like to acknowledge Prof. Subhajit Bandyopadhyay for providing the rotary-evaporator facility. Authors also express their deepest gratitude to Dr. Sumita Sengupta (nee Bandyopadhyay), University of Calcutta, for the mammalian expression vectors. The authors would also like to acknowledge Mr. Tamal Ghosh for their technical support in flow cytometry.

