## Supplementary File for "Association of miR-181a-5p with *Lantana camara* leaf extract-mediated inhibition of proliferation, survival, and migration in luminal A-type MCF-7 cells and triple-negative type MDA-MB-231 cells"

### Table S1. Details of the primer sequences used in the study

| **Target gene** | **Primer sequences** |
|---|---|
| 18S | Forward Primer: 5’-ACGGAAGGGCACCACCAGGAGT-3’<br>Reverse Primer: 5’-GAACGGCCATGCACCACCACC-3’ |
| Bcl-2 | Forward Primer: 5’-TCTTCAGGGACGGGGTGAACT-3’<br>Reverse Primer: 5’-TTCCACAAAGGCATCCCAGCC-3’ |
| Mcl-1 | Forward Primer: 5’-CAGAAAGAGGTGAGCTGTGTTAAAC-3’<br>Reverse Primer: 5’-CTAGCAAAGATGACCTTATGGCTCT-3’ |
| CDKN3 | Forward Primer: 5’-GAGTTTCGGGACAAATTAGCTGC-3’<br>Reverse Primer: 5’-CCCCGGAATATCTGCACATGTAC-3’ |
| U6 RT primer | 5’-CGCTTCACGAATTTGGCGTGTCA |
| miR181a SLP | 5’- GAAAGAAGGCGAGGAGCAGATCGAGGAAGAAGACGGAAGAAT GTGCGTCTCGCCTTCTTTCACTCACCG -3’ |
| U6 | Forward Primer: 5’- GCTTCGGCAGCACATATACTAAAAT -3’<br>Reverse Primer: 5’- CGCTTCACGAATTTGCGTGTCAT -3’ |
| miR181a | Forward Primer: 5’- GCAACATTCAACGCTGTCGG -3’ |
| qPCR reverse primer | 5’- CGAGGAAGAAGACGGAAGAA -3’ |
| CDKN3 primers used cloning | Forward Primer: 5’-TAAGCAAAGCTTGAGTTTCGGGACAAATTAGCTGC-3’<br>Reverse Primer: 5’- TGCTTAGGTACCGGAATATCTGCACATGTAC-3’ |
| CDKN3 primers used for SDM | Forward Primer: 5’-TGTTATCAACTTG**CCG**GTAAATGTACATGT-3’<br>Reverse Primer: 5’-ACATGTACATTTAC**CGG**CAAGTTGATAACA-3 |

**Table S2: Relative distribution of transiently transfected MCF-7 cells in different phases of the cell cycle under different treatment conditions with *Lantana camara* extract**

| Treatment conditions | MCF-7_Empty Vector | | | | MCF-7_pCmiR181a | | | |
|---|---|---|---|---|---|---|---|---|
| | Sub-G1 | G0/G1 | S | G2/M | Sub-G1 | G0/G1 | S | G2/M |
| Control | 8.77 ± 1.82 | 52.98 ± 1.53 | 15.98 ± 0.68 | 20.37 ± 2.08 | 16.33 ± 2.58 | 51.83 ± 1.69 | 16.16 ± 0.17 | 12.65 ± 3.37 |
| Vehicle control | 8.1 ± 2.52 | 56.3 ± 2.38 | 15.05 ± 1.15 | 18.87 ± 4.01 | 15.48 ± 2.94 | 55.93 ± 0.78 | 14.22 ± 0.31 | 12.38 ± 3.01 |
| 20 µg/mL | 16.85 ± 5.58 | 51.12 ± 1.23 | 13.43 ± 1.22 | 16.4 ± 2.59 | 41.75 ± 6.35 | 38.05 ± 3.5 | 9.98 ± 1.6 | 9.45 ± 2.19 |
| 40 µg/mL | 21.05 ± 8.83 | 45.07 ± 4.11 | 13.97 ± 1.2 | 15.65 ± 3.34 | 47.86 ± 2.85 | 31.85 ± 2.11 | 9.65 ± 1.16 | 9.51 ± 1.07 |
| 80 µg/mL | 27.97 ± 8.19 | 37.72 ± 3.44 | 12.3 ± 0.63 | 15.11 ± 3.14 | 58.3 ± 2.78 | 24.67 ± 1.43 | 8.48 ± 0.99 | 7.43 ± 1.48 |
| 120 µg/mL | 22.7 ± 9.51 | 41.42 ± 3.89 | 12.75 ± 0.61 | 15.93 ± 2.48 | 66.05 ± 8.14 | 20. 28 ± 5.03 | 7.25 ± 2.47 | 4.95 ± 1.85 |

**Table S3: Relative distribution of transiently transfected MDA-MB-231 cells in different phases of the cell cycle under different treatment conditions with *Lantana camara* extract**

| Treatment conditions | MDA-MB-231_Empty Vector | | | | MDA-MB-231_pCmiR181a | | | |
|---|---|---|---|---|---|---|---|---|
| | Sub-G1 | G0/G1 | S | G2/M | Sub-G1 | G0/G1 | S | G2/M |
| Control | 10.68 ± 4.17 | 52.47 ± 2.4 | 16.18 ± 0.54 | 20.9 ± 2.16 | 20.08 ± 1.67 | 51.6 ± 2.03 | 14.51 ± 1.02 | 13.75 ± 2.07 |
| Vehicle control | 8.45 ± 3.10 | 55.1 ± 2.5 | 16.05 ± 0.33 | 20.28 ± 1.19 | 22.88 ± 1.67 | 48.91 ± 1.8 | 14.1 ± 1.30 | 14.11 ± 1.78 |
| 40 µg/mL | 10.38 ± 3.51 | 55.61 ± 2.65 | 14. 85 ± 0.4 | 19.13 ± 1.32 | 19.9 ± 1.17 | 53.01 ± 2.14 | 13.21 ± 1.18 | 13.81 ± 1.69 |
| 60 µg/mL | 10.67 ± 3.58 | 56.5 ± 2.26 | 14.9 ± 0.53 | 18.45 ± 1.39 | 21.23 ± 1.62 | 52.28 ± 2.31 | 12.76 ± 1.26 | 13.6 ± 1.54 |
| 120 µg/mL | 11.95 ± 4.07 | 56.81 ± 0.91 | 11.82 ± 0.83 | 19. 37 ± 2.81 | 26.83 ± 3.16 | 49.67 ± 2.42 | 9.85 ± 1.38 | 13.61 ± 1.12 |
| 180 µg/mL | 14.35 ± 4.42 | 53.6 ± 1.85 | 10.87 ± 0.57 | 21. 18 ± 3.05 | 31.25 ± 1.05 | 44.48 ± 1.72 | 8.71 ± 0.43 | 15.56 ± 1.25 |

**Table S4: Comparative analysis of the dead cell population of transiently transfected MCF-7 cells under different treatment conditions with *Lantana camara* extract**

| Treatment conditions | MCF-7_Empty Vector | | | MCF-7_pCmiR181a | | |
|---|---|---|---|---|---|---|
| | Early apoptotic | Late apoptotic | Non apoptotic | Early apoptotic | Late apoptotic | Non apoptotic |
| **Control** | 2.28 ± 0.40 | 6.33 ± 2.53 | 7.81 ± 1.67 | 5.01 ± o.54 | 19.23 ± 3.35 | 13.93 ± 0.99 |
| **Vehicle control** | 3.8 ± 0.52 | 10.12 ± 3.32 | 7.73 ± 1.59 | 3.42 ± 0.42 | 21.68 ± 1.74 | 17.28 ± 1.22 |
| **40 µg/mL** | 3.51 ± 0.49 | 28.11 ± 1.29 | 14.25 ± 2.27 | 4.45 ± 0.86 | 29.51 ± 4.02 | 17.7 ± 3.15 |
| **80 µg/mL** | 4.68 ± 1.21 | 20.03 ± 1.71 | 12.98 ± 1.78 | 5.38 ± 1.21 | 26 ± 3.09 | 19.52 ± 3.09 |
| **120 µg/mL** | 3.36 ± 1.17 | 20.83 ± 1.41 | 24.51 ± 6.95 | 5.13 ± 1.96 | 28.88 ± 2.43 | 20.53 ± 4.05 |

**Table S5: Comparative analysis of the dead cell population of transiently transfected MDA-MB-231 cells under different treatment conditions with *Lantana camara* extract**

| Treatment conditions | MDA-MB-231_Empty Vector | | | MDA-MB-231_pCmiR181a | | |
|---|---|---|---|---|---|---|
| | Early apoptotic | Late apoptotic | Non apoptotic | Early apoptotic | Late apoptotic | Non apoptotic |
| **Control** | 3.84 ± 0.59 | 6.79 ± 0.76 | 2.5 ± 0.18 | 4.01 ± 0.34 | 9.37 ± 0.35 | 13.1 ± 1.84 |
| **Vehicle control** | 3.5 ± 0.71 | 5.8 ± 0.66 | 2.87 ± 0.26 | 4.4 ± 0.69 | 12.93 ± 0.92 | 14.98 ± 2.73 |
| **60 µg/mL** | 6.56 ± 2.03 | 10.5 ± 0.56 | 4.29 ± 0.37 | 2.68 ± 0.87 | 12.93 ± 1.31 | 32.46 ± 8.15 |
| **120 µg/mL** | 2.88 ± 0.90 | 17.82 ± 2.76 | 9.26 ± 1.3 | 0.67 ± 0.15 | 12.87 ± 3.56 | 39.81 ± 7.02 |
| **180 µg/mL** | 1.9 ± 0.55 | 24.94 ± 5.34 | 21.5 ± 2.90 | 1.12 ± 0.46 | 26.08 ± 12.21 | 28.82 ± 4.01 |
